# Whole genome sequencing of cell-free DNA yields genome-wide read distribution patterns to track tissue of origin in cancer patients

**DOI:** 10.1101/772657

**Authors:** Han Liang, Fuqiang Li, Sitan Qiao, Xinlan Zhou, Guoyun Xie, Xin Zhao, Kui Wu

**Author notes:** These authors contributed equally to this work (Han Liang, Fuqiang Li). To whom correspondence should be addressed (K.W.); (F.Q.L.).

## Abstract

Somatic mosaicism is widespread among tissues and could indicate distinct tissue origins of circulating cell-free DNA (cfDNA), DNA fragments released by lytic cells into the blood. By investigating the alignment patterns of whole genome sequencing reads with the genomic DNA of different tissues, we found that the read distributions formed type-specific patterns in some regions as a result of somatic mosaicism. We then utilized this information to construct a tissue-of-origin mapping model and evaluated its predictive performance on whole genome sequencing data from tissue and cfDNA samples. In total, 1,545 tissue samples associated with 13 cancer types were included, and identification of the tissue of origin achieved a specificity of 82% and a sensitivity of 80%. Furthermore, a total of 30 cfDNA samples from lung cancer and liver cancer patients and healthy controls were analyzed to predict their tissues of origin with a specificity of 87% and a sensitivity of 87%. Our results show that read distribution patterns from whole genome sequencing could be used to identify cfDNA tissues of origin with high accuracy, suggesting the potential application of our model to early cancer detection and diagnosis.

## Main

Somatic mosaicism is widespread among tissues (*1, 2*) and could indicate distinct origins for circulating cell-free DNA (cfDNA) fragments, which are released by lytic cells in the tissues into the blood plasma. Two groups have previously reported abnormal copy number variations (CNVs) in the cfDNA of pregnant women with cancers (*3, 4*). However, the researchers could not predict the cancer types based on the low-resolution CNV data when using calling algorithms with low-coverage whole genome sequencing. Here, we developed a sensitive model to directly identify somatic mosaicism footprints in the read distribution patterns of whole genome sequencing data.

To develop our model for tracking the tissue of origin of circulating tumor DNA (ctDNA), we first investigated the alignment patterns of sequencing reads from whole genome sequencing with the genomic DNA of 1,545 tissue samples associated with 13 cancer types from the Pan-Cancer Analysis of Whole Genomes (PCAWG) project of the International Cancer Genome Consortium (ICGC) and The Cancer Genome Atlas (TCGA) (*5, 6*). Each cancer type included more than 60 donors (**Table 1**). Our technology includes the following 4 major steps (**Methods**): 1) Compute the number of reads aligned with each fixed-width window. We divide each reference into a series of fixed-width windows; the typical window length is 10 kbp, an empirically obtained value. For simplicity, we join all the chromosomes together (Y excluded) and obtain a chain of 257973 windows (existing windows spanning two adjacent chromosomes). Then, we count the reads mapped inside each window for each sample to obtain the number of reads (NR). 2) Search for frequently occurring read distribution patterns among the samples. This step attempts to summarize the landscape of samples of the same type based on frequently occurring patterns, where a pattern refers to the relationships among the windows according to their NRs (higher/equal/lower) (**Figure 1a**). We considered that only windows in close proximity would influence each other effectively. Notably, one pattern could include several windows if their relationships are common among samples. As an example, consider the pattern “D, C, A, B”, in which the letters indicate the window indexes. This pattern means that for many samples, the NR of window D is higher than that of window C, which is higher than that of window A, which is higher than that of window B. In other words, we rank windows by their NRs to describe their relationships simply (**Figure 1b**). 3) Extract type-specific patterns from frequently occurring patterns. The previous step yields a large number of frequently occurring patterns by type. A pattern found often in one type of sample could also occur frequently in other types, and we need to extract those type-specific patterns to construct a model. Here, we transform the Fisher’s exact test *p*-value to measure how “specific” a pattern is for one type compared to another. The transformed value is used as the weight of pattern, and more highly significant *p*-values are always associated with higher weight values. Then, we extract patterns with weights above a calculated threshold. Obviously, when describing a type-specific pattern, we must note from which type the pattern is frequently and from which type the pattern is rarely. 4) Identify the type of sample according to the type-specific patterns. Two types of samples generate a paired type-specific pattern sets that are extracted together. Here, a pattern “match” a sample if the NR relationship among the windows described by the pattern is also valid for the sample. When we try to determine the possible type of a type-unknown sample based on these two types, we observe how many patterns from each type-specific pattern set match the sample. For each type, we accumulate the weights of the sample-matching patterns to calculate a score. The type-unknown sample will have two scores, one for each of the two types, and the type with the higher score would be considered the possible type of the sample. Obviously, if we have three or more types, we need to repeat step 3 for each pairwise combination of types and integrate all the results to obtain a final answer. A detailed schematic of the overall process is shown in **Figure 2**. After employing this method, we executed 5-fold cross validation on the tissue samples and found that our model achieved a high specificity of 82% and a high sensitivity of 80% (**Figure 3a**).

**Figure 1.**
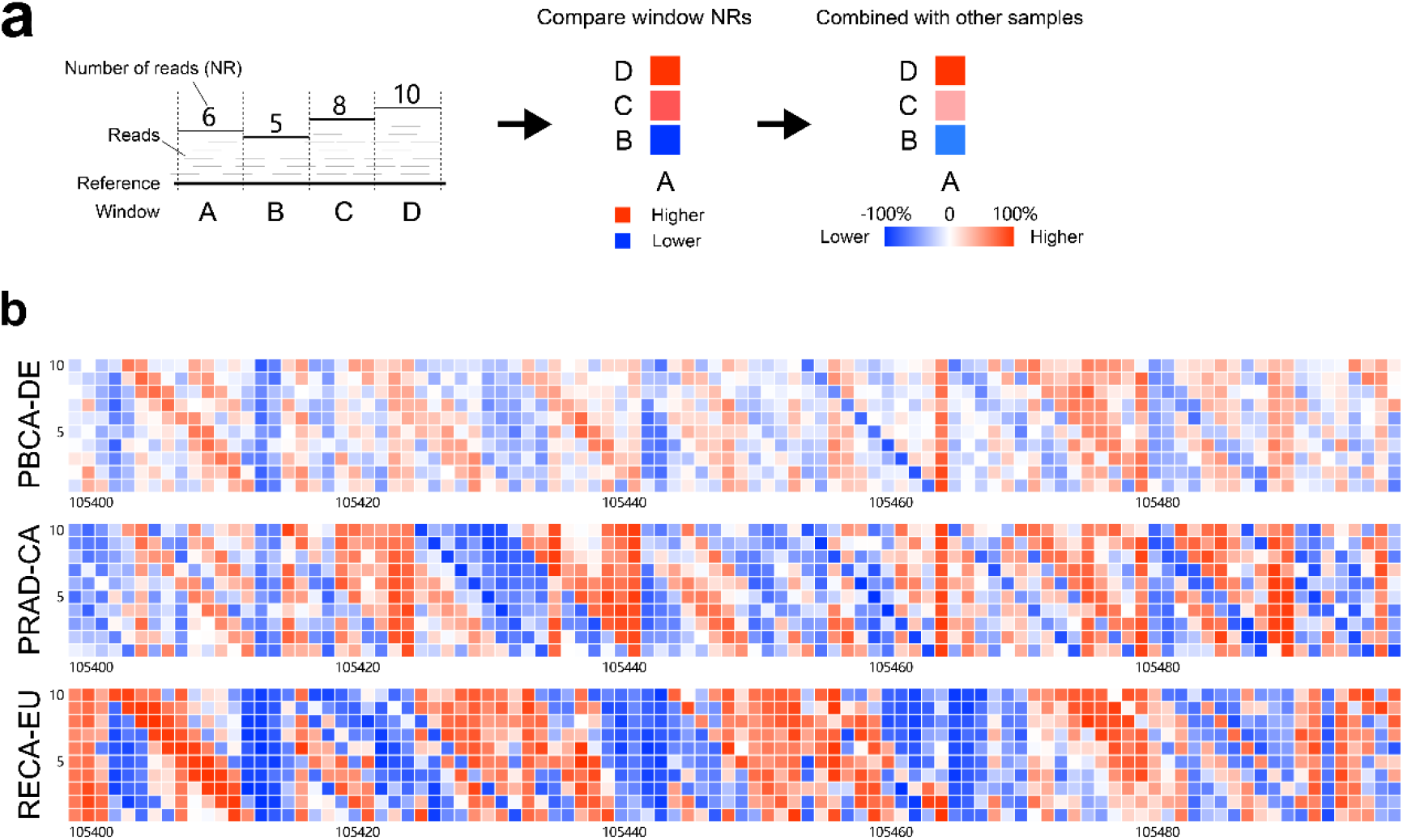
The read distribution pattern of a reference. a) The generation of read distribution patterns. As shown, we divided the reference into 4 fixed-width windows labeled A-D, counted the number of reads (NR) mapped to each window, and performed pairwise comparisons of the windows based on their NRs to obtain their relationships (higher/equal/lower). In this example, the NR of window A is lower than that of window B, indicated by a blue square, and higher than those of C and D, indicated by red squares. To discuss the relationships between two windows in the context of multiple samples, we can assume that the two windows are labeled A and B. Then, we use a percentage to represent the difference between samples for which the NR of A is higher than that of B (n=Na) and those for which the NR of B is higher than that of A (n=Nb): the percentage value=(Na-Nb)/(Na+Nb)*100%. b) The read distribution patterns of three types of tissue samples. We joined all the chromosomes (Y excluded) together and obtained a long chain of 257973 windows (for hg19). The relationships among the windows ranked 105400-105500, 100 windows in total, are shown for each group. Each window was compared with its 10 downstream windows. The x-axis represents the window index, the y-axis represents the distance of the downstream windows from the current window, and the colors represent the percentages calculated with the method described in a).

**Figure 2.**
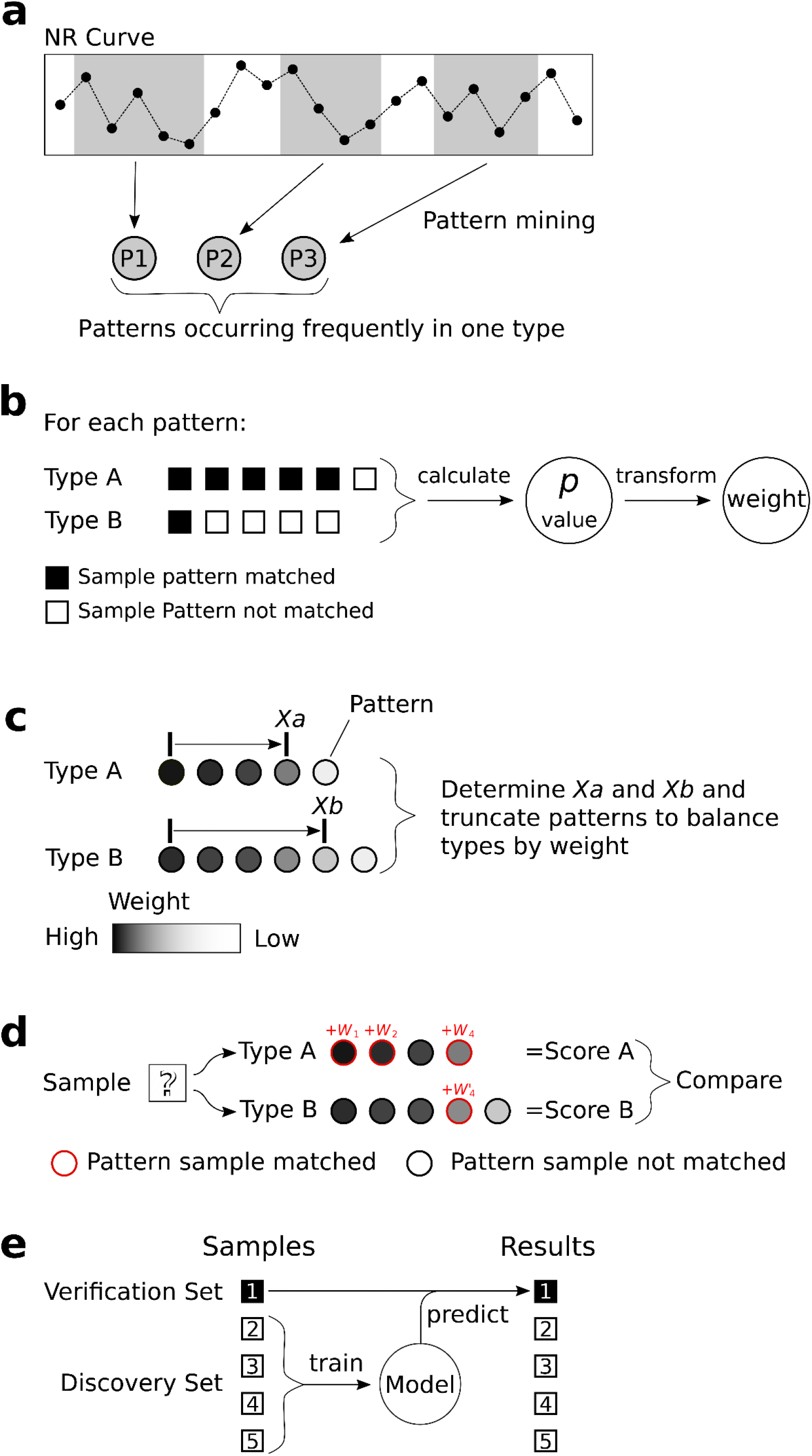
Schematic of model construction and prediction. a) Pattern mining. We searched for frequently occurring patterns in the NR curve for each type of sample. A pattern is considered frequent if its shape (the order of windows ranked by their NRs) is the same for most (e.g., 60%) of the samples’ NR curves. 2) Type-specific pattern extraction. For each pattern, we calculated the Fisher’s exact test *p*-value by checking how many samples were matched by the pattern in each type and then transformed the *p*-value to derive the weight of pattern. We applied these operations to each pattern and retained patterns with a *p*-value≤0.01 as type-specific patterns. 3) Pattern balancing. We mined for frequently occurring patterns in type A samples, excluded the patterns found in type B samples and then repeated these steps by exchanging types A and B to obtain two type-specific pattern sets. However, the resulting pattern sets were not necessarily balanced in weight. To resolve this problem, we used an optimization algorithm to derive *Xa* and *Xb* and retained the top *Xa* highest-weight patterns for type A and the top *Xb* for type B to balance the weights. d) Sample prediction. To determine the type of a type-unknown sample based on two type-specific pattern sets, we checked whether the patterns matched the sample and summed the weights of the sample-matched patterns for each type to obtain two scores, one per possible type. The type with the higher score would be considered the sample’s possible type. e) We divided the tissue samples equally into 5 subsets and the plasma samples into 10 subsets. For each subset, we combined the remaining subsets into a discovery set to train a model and then predicted the original subset as the verification set. Then, we combined all the predicted results to analyze the specificity and sensitivity of the model.

**Figure 3.**
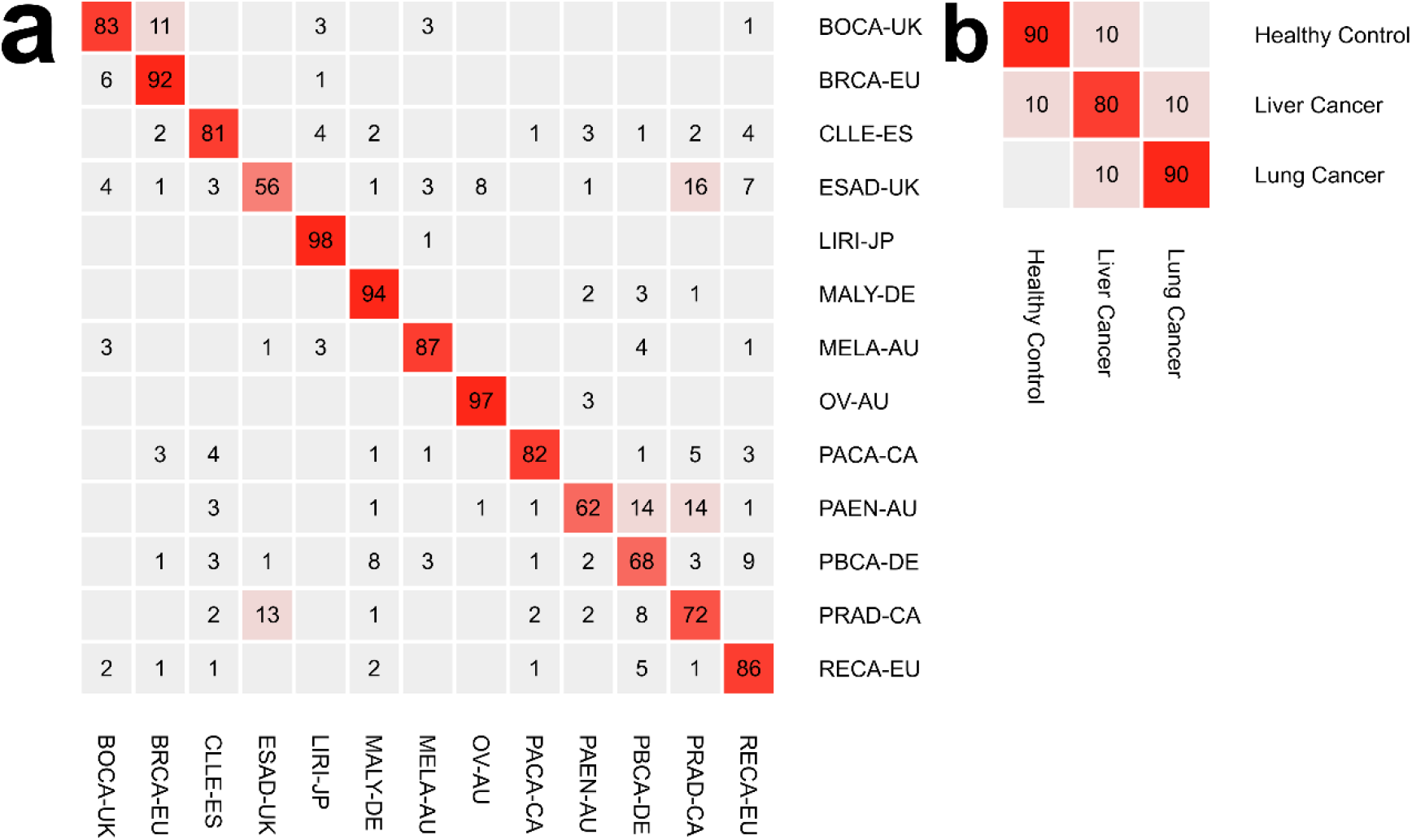
The predicted results of the tissue and cfDNA samples. a) The results of the tissue samples. This test involved 13 types of tissue samples, and the figure shows the integration of the 5-fold cross validation results. The rows represent the different types of samples, the y-axis labels (at the right) represent the real sample types, the columns represent the predicted results, and the x-axis labels (at the bottom) represent the predicted types; the numbers inside the cells represent the percentages of samples predicted as each x-axis label among the samples marked with each y-axis label. b) The results of the cfDNA samples. This test involved 3 types of cfDNA samples, and the figure shows the integration of the 10-fold cross validation results.

**Table 1.** Details of Cancer Types from the PCAWG Project

| Cancer Type from the PCAWG Project | Cancer Type Abbreviation | Number of Donors |
| --- | --- | --- |
| Bone Cancer - United Kingdom | BOCA-UK | 76 |
| Breast ER+ and HER2- Cancer - European Union/United Kingdom | BRCA-EU | 79 |
| Chronic Lymphocytic Leukemia - Spain | CLLE-ES | 100 |
| Esophageal Adenocarcinoma - United Kingdom | ESAD-UK | 100 |
| Liver Cancer - Japan | LIRI-JP | 259 |
| Malignant Lymphoma - Germany | MALY-DE | 101 |
| Skin Cancer - Australia | MELA-AU | 70 |
| Ovarian Cancer - Australia | OV-AU | 73 |
| Pancreatic Cancer - Canada | PACA-CA | 148 |
| Pancreatic Cancer Endocrine Neoplasms - Australia | PAEN-AU | 69 |
| Pediatric Brain Cancer - Germany | PBCA-DE | 251 |
| Prostate Adenocarcinoma - Canada | PRAD-CA | 124 |
| Renal Cell Cancer - European Union/France | RECA-EU | 95 |

To evaluate our model’s identification of tissues of origin for cfDNA samples, a total of 30 cfDNA samples from lung cancer and liver cancer patients and healthy controls were analyzed. The cfDNA samples were sequenced on a BGISEQ-500 with an average 3X depth of coverage. To explore the samples sufficiently, we performed 10-fold cross validation on the cfDNA samples (rather than 5-fold, which was used for the tissue samples, as 13 types of tissue samples had to be distinguished and the cost of 10-fold cross validation would have been excessive). We found that our model achieved a high specificity of 87% and a high sensitivity of 87% (**Figure 3b**). There were 4 erroneous results among the 30 samples: 1 healthy control sample was misidentified as liver cancer, 1 lung cancer sample was misidentified as liver cancer, and 2 liver cancer samples were misidentified, one as healthy control and one as lung cancer. Notably, our model distinguished healthy control samples with high accuracy, which is very important in early tumor screening.

We consider that CNV data are not always sensitive enough to describe cfDNA features given the low tumor cell DNA concentration in cfDNA. The type-specific patterns of read distribution used in our model are based on the NR relationships among windows. In theory, the NR relationship between two windows from the same sample is not affected by the sequencing depth, and in practice, this method works quite well with a low sequencing depth. However, the biological meaning of read distribution patterns is still ambiguous and needs more exploration. Understanding the biological significance of these patterns would help us more effectively improve our method and support the discovery of cancer mechanisms and new cancer treatments.

## Author contributions

K.W. and F.Q.L. conceived of the idea and supervised the work. H.L. developed the model of tracking tissue-of-origin for ctDNA. S.T.Q. and G.Y.X. performed standard pipeline of sequencing data. X.L.Z. and X.Z. performed experiments of sequencing. H.L., F.Q.L wrote the manuscript. K.W. contributed to drafting and revising the manuscript.

## Data and materials availability

Alignment files of 1,545 tissue samples involving 13 cancer types from the Pan-Cancer Analysis of Whole Genomes (PCAWG) project were analyzed on the Cancer Genome Collaboratory, an academic compute cloud resource that allows researchers to run complex analysis operations across large ICGC cancer genome data sets. The sequencing data of 30 cfDNA samples have been deposited in the CNSA (https://db.cngb.org/cnsa/) of CNGBdb with accession code CNP0000680. The sequencing data of cfDNA samples are available on reasonable request. An up-to-date version of the analysis code, along with an up-to-date README, will available as a Github repository.

We are thankful to the production team of China National GeneBank, Shenzhen, China.

## Methods

### Counting the read distribution on the reference

Here, we define sorted window indexes (SWIs), which refer to a series of indexes of windows placed on the reference. These indexes are sorted according to the number of reads mapped inside each window, or NR. In this paper, an SWI is considered a simplified read distribution.

First, we need to determine every sample’s SWI. This step is performed as follows:

1. Divide the reference into a series of fixed-width windows, and label those windows with their indexes. For simplicity, we join all the chromosomes together in numerical order (chromosomes 1-22 & X; Y excluded). The window length is typically set as 10 kbp, an empirical value, but sometimes, another value in the range of 5 kbp-50 kbp is used.
2. Count the reads mapped inside each window for each sample. The reads are mapped to the reference with a standard short read alignment method. When a read spans two windows, we consider the window in which most of its bases are located its mapped window.
3. Obtain the SWI for each sample by sorting the window indexes by NR.
4. Repeat steps 2 and 3 until all samples have been processed.

Each sample produces an SWI, and the SWIs gained from a sample set form an SWI set.

### Search for frequent distribution patterns among SWIs

A pattern is a series of order-sensitive numbers that refer to window indexes, e.g., (3, 1, 2); it is a miniature SWI. A pattern could contain a series of numbers too long to search directly, and we developed a splicing model to find such patterns:

If there are two patterns, one of which has a tail section identical to another pattern’s head section, the operation of joining the former pattern with the latter pattern is called splicing.

As an example, consider two patterns, (1, 2) and (2, 3). The tail section of the former pattern is “2”, which is the same as the latter pattern’s head section. We take the former pattern (1, 2) and splice on the latter pattern’s remaining tail section (3) to gain a new, longer pattern (1, 2, 3).

To reduce computational complexity, however, we splice only pairs of patterns whose indexes contain only one different element. For example, we could splice patterns (1, 2, 3) and (2, 3, 4) into a new pattern (1, 2, 3, 4), because the former and the latter contain only one different element each (“1” and “4”, respectively).

Splicing combines two shorter patterns to produce a longer pattern, and the shortest pattern contains only two indexes and is called an L2. We searched the L2s using the following formula:

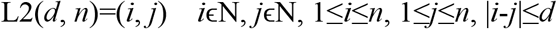

where *n* is the maximum index among the windows, and *d* is the maximum distance between two indexes of L2. Based on empirical evidence, we set *d* as 40.

We are interested in frequently occurring patterns, and therefore, additional checks are required. To determine whether a pattern occurs frequently, we determine how many SWIs a pattern match. Here, we say a pattern “match” an SWI when the order of the windows contained in the pattern is the same as that in the SWI. For example, we say the pattern (3, 1, 2) matches the SWI (4, 3, 1, 2) because the order of “1”, “2”, and “3” in the pattern is the same as that in the SWI. Furthermore, the number of samples matched by a pattern must reach a given threshold to be considered a frequent pattern. Then, we attempt to obtain all frequent patterns by splicing the patterns iteratively until no more frequent patterns are generated.

Finally, the frequent patterns obtained from a sample set form a frequent pattern set.

### Extracting type-specific patterns from frequent patterns

A frequent pattern is type-specific if there is a significant difference in its coverage between two sample sets. First, we filter frequent patterns according to their coverage of two sample sets. Two pattern sets gained from two sample sets, after each is filtered with the other sample set, will generate two paired type-specific pattern sets. To measure the ability of a frequent pattern to distinguish two kinds of samples, we use the transformed Fisher’s exact test *p*-value as the pattern’s weight. To calculate the *p*-value, we need to check how many samples in each sample set are matched by the pattern. The transformation formula is

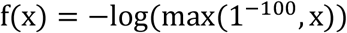

To assign a sample to one of the two types, we use the following formula:

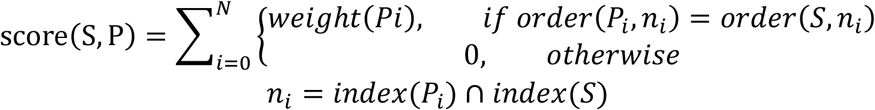

where S is an SWI extracted from the type-unknown sample, P is one of the two paired type-specific pattern sets, N is the total number of patterns in P, weight(x) is the weight of pattern x, Pi is the *i*-th pattern in P, order(x, y) is the order vector of index set y in pattern/SWI x, and index(x) is the index set of pattern/SWI x.

Comparing the scores of the two type-specific pattern sets, the type corresponding to the higher score will be considered the possible type.

However, the differences between the weighted sums of two specific pattern sets can be very large, making one score always higher than another. Thus, we need to delete some patterns to balance the weights, which is done with the following formula:

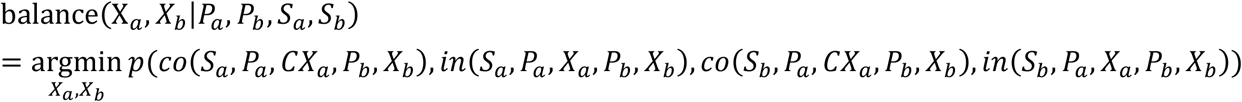

where X_a_ and X_b_ are the two factors to be determined. X_a_ is used as a length to retain the top *X_a_* highest-weight patterns of the type-specific pattern set P_a_, which was obtained from sample set S_a_; X_b_, P_b_ and S_b_ belong to the other sample type and have the same meanings as X_a_, P_a_ and S_a_, respectively. p(*a, b, c, d*) is the *p*-value calculated using Fisher’s exact test with the factors *a, b, c, d*; co(*s, a, b, c, d*) is the number of samples predicted correctly in sample set s with pattern set *a* truncated with length *b* and pattern set *c* truncated with length *d*; in(*s, a, b, c, d*) is the number of samples predicted incorrectly with the same factors as function co.

To solve this formula, we use an iterative algorithm. First, we set X_a_ as a reasonable random value and find the best X_b_ in the given situation; then, we keep X_b_ unchanged and find the best X_a_. The end condition of this iterative process is stable values of X_a_ and X_b_.

Finally, we use X_a_ and X_b_ to truncate the two paired type-specific pattern sets.

### Identify a sample’s type according to type-specific patterns

The method used to identify a sample’s type was introduced in the previous step for only two types. When trying to determine a sample’s type from more than two candidates, we need to repeat the previous step for every combination of two types. Obviously, for N types, there will be N(N-1)/2 combinations. In this situation, every repetition will provide a possible answer, and all these answers can be combined to determine a final prediction.

